# Uncertainty Quantification in Stochastic Dynamical Gene Regulatory Networks

**DOI:** 10.64898/2026.08.31.747806

**Authors:** Francisca Pizarro Galleguillos, Satyajeet Bhonsale, Jan F.M. Van Impe

## Abstract

The dynamics of gene regulatory networks are governed by intrinsic noise, stemming from the random nature of biochemical reactions, and by extrinsic noise, arising from fluctuations in cellular components and environmental conditions. Together, these sources can compromise the reliability of predictive computational models if not properly accounted for, and capturing both effects within a single framework remains a non-trivial task in computational biology. In this work, we propose an uncertainty quantification framework that addresses these two contributions jointly: intrinsic stochasticity is described through a partial integro-differential equation (PIDE) for the protein probability density function, whereas extrinsic noise is represented as parametric uncertainty in the kinetic parameters. The propagation of the uncertainty is carried out via an intrusive polynomial chaos expansion (PCE), in which the PCE coefficients are obtained from a stochastic Galerkin projection of the PIDE, yielding a coupled deterministic system that is solved with standard numerical methods. We illustrate the approach on a positive autoregulatory gene network with one and two uncertain kinetic parameters. The proposed approach accurately reproduces the mean, variance, and full protein probability density function, including the bimodal distributions, at a substantially lower computational cost.

## 1 Introduction

A gene regulatory network (GRN) consists of a collection of genes, proteins, and other molecules that interact to regulate gene expression within a cell [1]. Fundamentally, these networks are constructed from the interplay of two elements: genes that encode transcription factors, and regulatory modules on the DNA, such as promoters and enhancers, which modulate transcriptional activity [2]. GRNs play a central role in diverse cellular processes, including metabolism, cell-cycle control, and responses to environmental signals [3], [4]. Recent advances in high-throughput and singlecell technologies have provided unprecedented insight into the structure and dynamics of GRNs [5]. These advances highlight the need for modelling approaches capable of interpreting increasingly detailed experimental observations, making predictive computational models indispensable for analyzing network functionality and complex regulatory mechanisms [1].

Gene expression is inherently stochastic [6], [7], influenced by intrinsic noise originating from the random timing of biochemical reactions, and extrinsic noise arising from fluctuations in cellular components and environmental conditions [3]. When molecular components are abundant, deterministic models utilizing ordinary differential equations (ODEs) and mass-action kinetics can characterize the average temporal behavior of gene products [8]. Although computationally efficient and valuable for understanding system dynamics, these models fail to capture the inherent randomness observed in gene expression [9]. Stochastic models address this limitation by accounting for the probabilistic nature of molecular interactions, thereby representing intrinsic noise, particularly in contexts with low molecular counts [10], [11].

A principled framework to describe this distributional dynamics is provided by the Chemical Master Equation (CME), which governs the time evolution of the probability distribution over molecular states. However, its analytical solution is often intractable. To overcome this, stochastic simulation algorithms (SSA), such as the Gillespie algorithm [12], [13], are widely used to generate exact sample paths of the stochastic process governed by the CME. Although effective, these methods are often computationally demanding, particularly for large-scale regulatory networks or when repeated simulations are required. This motivates the development of scalable distribution-based approximations.

Several such strategies have been proposed to address the computational complexity of the CME. Diffusion-based approaches, such as the chemical Fokker–Planck equation [14], as well as the van Kampen system-size expansion and its linear noise approximation (LNA) [15], approximate the dynamics of the CME by continuous stochastic processes and provide fast to evaluate estimates of the marginal distribution under near-Gaussian fluctuations and large molecule numbers (i.e. the thermodynamic limit). However, these approximations often lose accuracy in GRN characterized by low molecular copy numbers, nonlinearities, or multimodal distributions [16], [17]. Alternative methods, including finite state projection (FSP) [18] and moment-based approaches [19], [20], which reduce the computational burden by truncating the state space and evolving low-order moments, respectively. Normally they do not compute a marginal distribution, and still, suffer from scalability issues or closure inaccuracies in complex systems and low predictive capacity [21]. In these settings, multidimensional partial integro-differential equations (PIDEs) provide an accurate and scalable continuous representation of the CME, naturally accommodating regulatory gene expression dynamics [22], [23].

Regardless of the computational framework, GRN models are approximations of biological reality and therefore subject to multiple sources of uncertainty. These include incomplete knowledge of network topology and reaction mechanisms (model uncertainty) [24], imprecise parameter estimation (parametric uncertainty) [17], and the intrinsic stochasticity of molecular interactions (aleatory uncertainty). Such uncertainties can substantially affect model predictions and, if unaccounted for, may lead to misleading interpretations and unreliable conclusions in biological applications.

The systematic characterization and propagation of these uncertainties in model predictions is known as uncertainty quantification (UQ) [25]. In computational biology, UQ remains a non-trivial task due to the strong nonlinearity, high dimensionality, and substantial computational cost of many mechanistic models [26]. These challenges are particularly pronounced in gene regulatory networks, where nonlinear interactions can generate complex and often multimodal predictive distributions, even in relatively simple architectures [24], [27], [28]. While a broad range of UQ methodologies has been developed and applied in deterministic biological models [26], [29], [30], their extension to stochastic GRN models remains comparatively limited.

To address these limitations, surrogate modelling techniques are commonly employed in uncertainty quantification to reduce the computational cost associated with complex mechanistic models. These methods construct an approximate model or surrogate that is computationally cheaper to evaluate while retaining essential features of the original model [31]. Among these approaches, polynomial chaos expansion (PCE) [32], [33] has emerged as a fast and accurate framework for representing and propagating parametric uncertainty. Applications to biological systems have demonstrated its effectiveness in deterministic settings [34], [35], [36], [37]. However, their application to stochastic biological models, and in particular to stochastic gene regulatory networks, remains largely unexplored. Existing efforts have focused on inference-oriented approaches rather than on the propagation of parametric uncertainty through stochastic dynamics. For instance, [38] introduces a surrogate based on a finite-state Markov chain with negative binomial output distributions to approximate the CME from sampled GRN data. Similarly, [39] develops a framework to infer extrinsic noise in gene expression using hierarchical Markov models combined with sequential Monte Carlo techniques.

In this paper, we propose an uncertainty quantification framework for stochastic dynamical gene regulatory networks based on polynomial chaos expansions. The PCE coefficients are obtained using intrusive stochastic Galerkin projections applied to PIDE stochastic models. While Hill-type regulatory functions are used for demonstration, the framework can accommodates alternative kinetic forms through appropriate definitions of the reaction-rate expressions. We demonstrate the efficacy of the proposed method using a positive autoregulatory network as a case study, showing that it delivers accurate uncertainty estimates while remaining computationally tractable.

The paper is organized as follows: Section 2 details the principles of PCE, the mathematical modelling of stochastic GRNs, and their Galerkin approximation; Section 3 presents the results of applying our method to two case studies, and Section 4 concludes with a discussion of findings and future research directions.

## 2 Methods

### 2.1 Mathematical Model of Stochastic Dynamical Gene Regulatory Network

We consider a single-gene auto-regulatory network expressing a single protein type *X* through its messenger RNA, *mRNA*, following the central dogma. The number of mRNA and protein molecules is represented by *m, x* ∈ ℝ_+_, respectively. A schematic representation of the gene network is shown in Figure 1.

**Figure 1:**
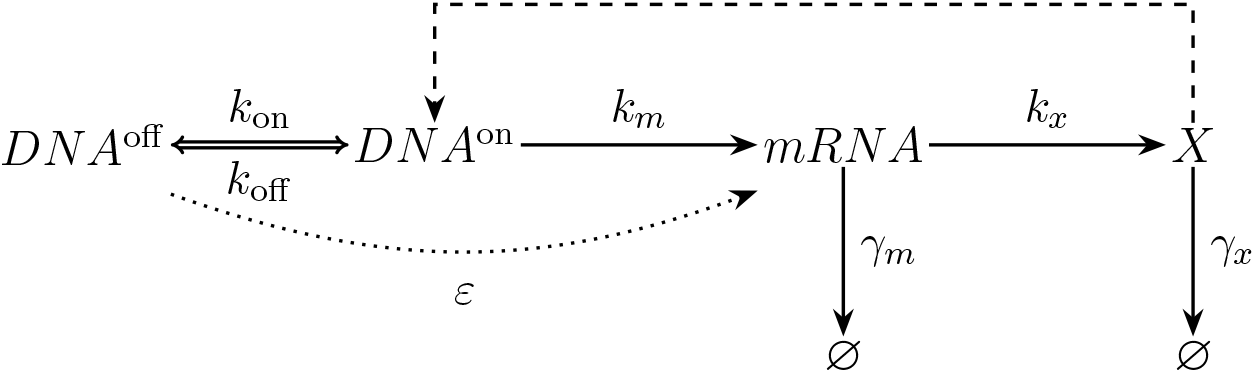
Schematic representation of the stochastic gene expression model regulated by protein *X*. The gene can switch between an active state (*DNA*^on^) and an inactive state (*DNA*^off^ ). In the active state, the gene transcribes mRNA at a rate *k*_*m*_, which is then translated into protein *X* at a rate *k*_*x*_. The gene can also switch back to the inactive state at a rate *k*_off_, while the reverse switching occurs at a rate *k*_on_. Additionally, there is a leakage rate *ε* allowing transcription from the inactive state. Both mRNA and protein degrade at rates *γ*_*m*_ and *γ*_*x*_, respectively. The regulation by protein *X* is indicated by the dashed line.

In the model, proteins are assumed to be produced in bursts. This implies that the degradation rate of mRNA is significantly higher than that of proteins (*γ*_*m*_ ≫ *γ*_*x*_). Based on this assumption, proteins can be considered to be produced in random uncorrelated events. The conditional probability for the protein level to jump from state *y* to state *x* is given by *β*(*x* − *y*), where the burst follows an exponential distribution [22]:

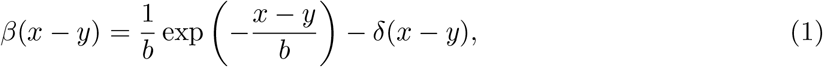

With 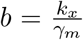 denoting the mean number of proteins produced per burst and *δ* the Dirac delta function.

Under the assumptions outlined above, the time evolution of the protein probability density function *p*(*t, x*) is governed by a partial integro-differential equation (PIDE), as first formulated by Friedman *et al*. [22] and later extended to its multivariate form by Pájaro *et al*. [23]. The PIDE model for the single gene regulatory network is given by:

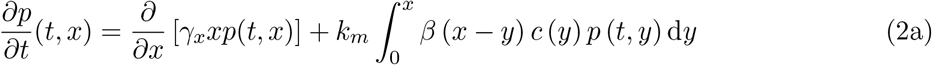

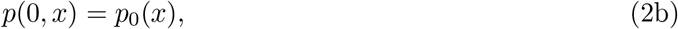

where *p* : ℝ_+_ ×ℝ_+_ → ℝ_+_ denotes the probability density function, **x** assumes continuous values in ℝ_+_ and represents a continuous approximation to the number of proteins described by the CME. The first term in the right-hand side of the equation represents protein degradation, and the integral describes protein production by bursts. The initial condition *p*_0_(**x**) is a probability density function assumed to follow a normal distribution.

The function *c*(*x*) represents the regulatory control of the gene by *X*, it is typically modelled using Hill functions to capture activation and repression mechanisms [23], [40]. The general form of the Hill function is given by:

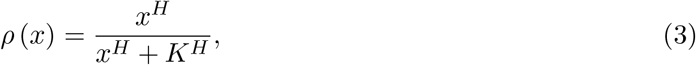

where *H* is the Hill coefficient indicating the cooperativity. A positive value indicates that *X* inhibits transcription, corresponding to negative feedback (repression), while a negative value indicates that *X* promotes transcription, corresponding to positive feedback (activation). The parameter *K* is the corresponding Hill constant. In the case of regulation by a single protein, the transcription rate is proportional to the fraction of active promoter, expressed as:

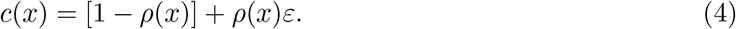

Here *ε* is the leakage constant from the inactive promoter. The effect of multiple proteins regulating a gene can be modelled by combining individual Hill functions using logical operations [40].

### 2.2 Polynomial Chaos Expansion

PCE is a spectral method for uncertainty quantification that represents random variables as expansions in orthogonal polynomial bases. Originally introduced by Wiener [32] and later extended by Ghanem and Spanos [33], PCE has become a widely used framework for uncertainty analysis in complex systems.

Let ***θ*** = [*θ*_1_, *θ*_2_, …, *θ*_*nθ*_ ]^*T*^ be a vector of *n*_*θ*_ independent standard random variables, and let *u* = *u*(***θ***) be a function of ***θ***. The PCE representation of *u* is expressed as:

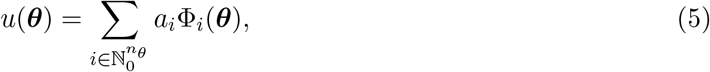

where *a*_*i*_ are the PCE coefficients and 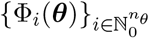 is a set of multivariate orthogonal polynomials, indexed by the multi-index 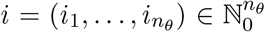, where *i*_*k*_ is the degree of the univariate polynomial in the *k*-th variable and 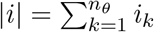 denotes its total degree. For any two functions *g* and *h* of ***θ***, we define the inner product ⟨*g h*⟩ as the expectation with respect to the joint probability density function *f*_***θ***_(***θ***) of ***θ***,

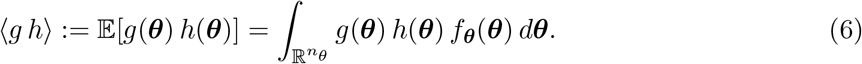

The polynomials 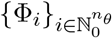 are orthogonal with respect to this inner product,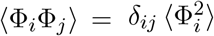, where *δ*_*ij*_ is the Kronecker delta and ⟨Φ^2^⟩ is the squared norm of Φ_*i*_. Each uncertain parameter can be expressed as a function of a random variable, and the Wiener–Askey scheme is then used to construct families of orthogonal polynomials associated with the probability distribution of the parameters [41].

To make the PCE representation computationally feasible, it is necessary to truncate the infinite series to a finite number of terms. A common approach is the total-degree truncation, which limits the total degree of the polynomials to a maximum value *d*. This defines the truncated multi-index 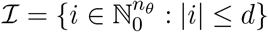. The truncated PCE can be expressed as:

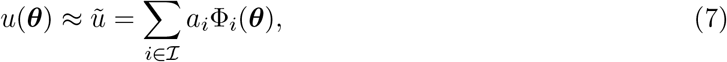

Where 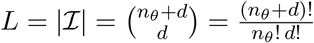 is the total number of polynomial terms.

There are two main approaches to calculate the PCE coefficients: the intrusive and the nonintrusive approach. In the non-intrusive approach, the model is treated as a black box from which samples are drawn, and the coefficients are obtained from the resulting model evaluations using regression, projection or stochastic collocation techniques [42]. In contrast, the intrusive approach reformulates the governing equations of the model by substituting the truncated PCE representation into them, yielding a set of deterministic equations for the PCE coefficients [43].

In this work we follow the intrusive approach, which relies on the Galerkin projection method [33]. The inner product defined in (6) is employed to project the PCE representation onto each basis polynomial Φ_*j*_:

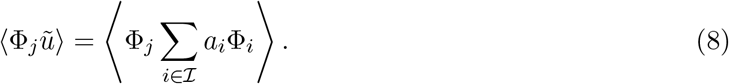

By the linearity of the inner product, and since the coefficients *a*_*i*_ do not depend on ***θ***, they can be taken outside the inner product, yielding:

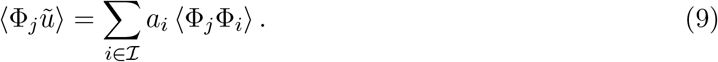

Using the orthogonality property of the polynomials, the expression simplifies to isolate the coefficients *a*_*j*_:

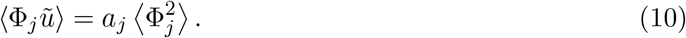

Finally, the PCE coefficients are given by:

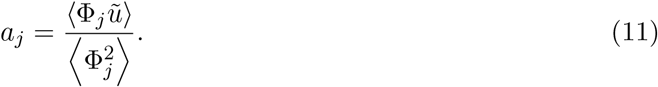

When this projection is applied to the governing equations of the stochastic dynamical system, it produces a coupled set of deterministic equations for the coefficients *a*_*j*_, which can be solved using standard numerical methods [43].

Once the PCE coefficients are available, the expected value and variance of the random variable *u* can be obtained directly from them as:

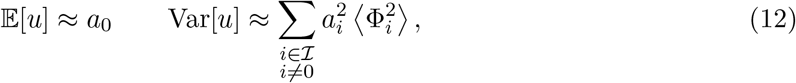

Where 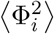 can be computed upfront and equals 1 for normalized polynomials. These expressions follow from the orthogonality of the polynomial basis and therefore hold regardless of how the coefficients *a*_*i*_ are computed.

### 2.3 Galerkin Approximations of PIDEs

The PIDE model introduced in Equation (2) depends on a set of kinetic parameters. In practice, these parameters are subject to uncertainty due to measurement errors or cell-to-cell variability. To account for this uncertainty, we model the kinetic parameters as random variables and apply the Galerkin method to derive a system of deterministic equations for the PCE coefficients of the solution. In what follows, we first derive the projected system for a single uncertain parameter, and then extend the procedure to two independent uncertain parameters. The same methodology generalizes to an arbitrary number of uncertain parameters.

#### 2.3.1 One-dimensional equation with uncertain protein degradation rate

Let us consider a one-dimensional PIDE defined on [0, *T* ]×[0, *ℓ*_*x*_] with uncertain protein degradation rate *γ*_*x*_, modelled as a real-valued random variable. The protein number domain is truncated at *x* = *ℓ*_*x*_ chosen large enough and a homogeneous Dirichlet boundary condition is imposed at the inflow boundary. The resulting equation reads:

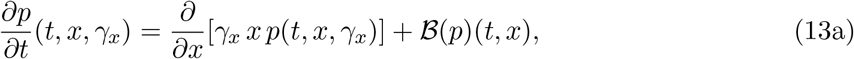

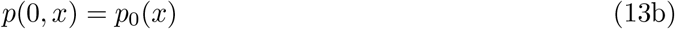

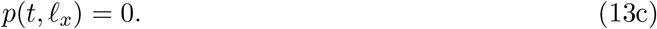

where B(*p*) denotes the burst-production integral operator introduced in Equation (2), which is linear in *p* and, in this case, does not depend on *γ*_*x*_. The uncertainty in *γ*_*x*_ propagates to the solution *p*(*t, x, γ*_*x*_), which we approximate using a truncated PCE of total order *d*:

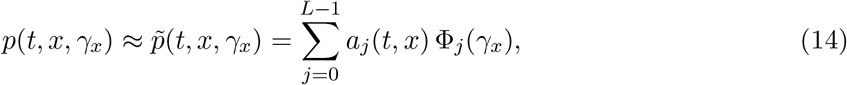

where 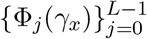 are orthonormal polynomials with respect to the probability density function of *γ*_*x*_, and *a*_*j*_(*t, x*) denote the corresponding PCE coefficients. The Galerkin projection is then carried out in two steps: substitution of the PCE expansion into the PIDE and projection onto each basis polynomial with respect to *γ*_*x*_.

Inserting the PCE approximation (14) into the PIDE (13) and exploiting the linearity of B yields

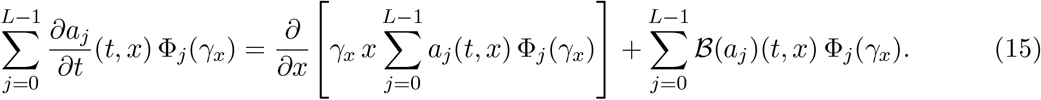

Multiplying (15) by an arbitrary basis function Φ_*k*_(*γ*_*x*_), with *k* ∈ {0, …, *L* − 1}, gives

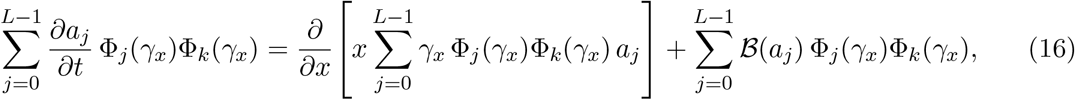

where the dependence on (*t, x*) has been omitted for brevity.

Finally, taking the expectation with respect to the distribution of *γ*_*x*_ on both sides of (16) and using the orthonormality property ⟨Φ_*j*_Φ_*k*_⟩ = *δ*_*jk*_, the time-derivative and burst terms collapse to the single coefficient *k*, while the degradation term retains a full coupling through the moments ⟨*γ*_*x*_ Φ_*j*_Φ_*k*_⟩. The resulting coupled system of deterministic PIDEs reads, for each *k* ∈ {0, …, *L* − 1},

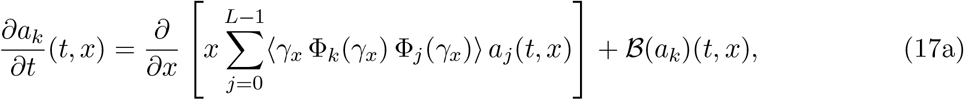

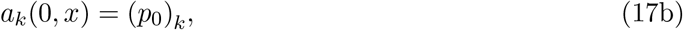

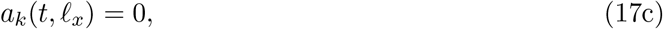

where (*p*_0_)_*k*_ denotes the *k*-th PCE coefficient of the initial condition. Since *p*_0_(*x*) is independent of *γ*_*x*_, only its zeroth coefficient is nonzero, i.e. (*p*_0_)_0_ = *p*_0_(*x*) and (*p*_0_)_*k*_ = 0 for *k* ≥ 1. The homogeneous Dirichlet boundary condition is preserved component-wise since all PCE coefficients of the zero function vanish. Since B does not depend on *γ*_*x*_, its projection onto Φ_*k*_ reduces to B(*a*_*k*_), and no additional coupling is introduced through the burst term in this case. For simplicity, we define the vector of PCE coefficients **a**^*L*^ := (*a*_0_, *a*_1_, …, *a*_*L−*1_)^*T*^. Then, Equation (17) can be rewritten in vector form as

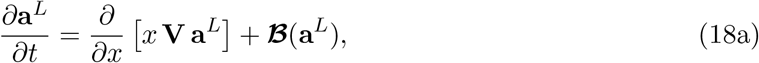

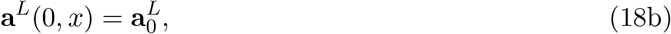

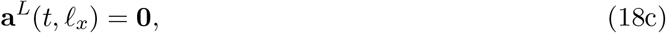

where ***B***(**a**^*L*^) denotes the component-wise application of B to the entries of **a**^*L*^, 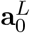 collects the PCE coefficients of the initial condition, and **V** ∈ R^*L×L*^ is the projection matrix with entries

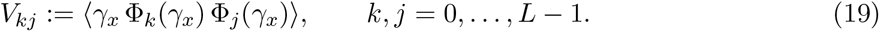

The entries of **V** depend only on the distribution of *γ*_*x*_ and on the chosen polynomial basis, and can therefore be precomputed using numerical quadrature or, for standard distributions, known closed-form expressions [44].

#### 2.3.2 One-dimensional equation with uncertain protein degradation rate and transcription rate

Consider the same one-dimensional PIDE on [0, *T* ] × [0, *ℓ*_*x*_] with two independent uncertain parameters: the protein degradation rate *γ*_*x*_ and transcription rate *k*_*m*_. We denote the joint random vector as ***θ*** = [*γ*_*x*_, *k*_*m*_]:

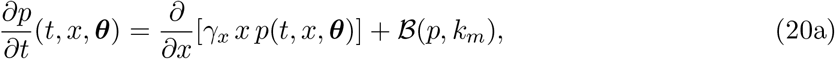

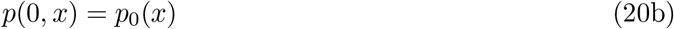

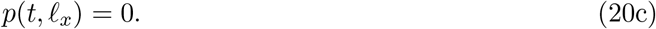

The burst-production operator is written as

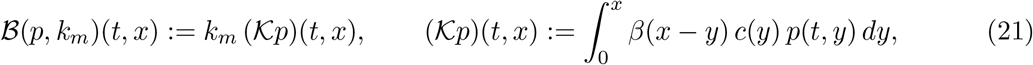

Since *γ*_*x*_ and *k*_*m*_ are independent, we construct a tensor-product PCE basis

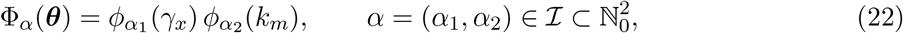

where {*ϕ*_α1_ (*γ*_*x*_)} and {*ϕ*_α2_ (*k*_*m*_)} are the univariate orthonormal polynomials associated with the marginal distributions of *γ*_*x*_ and *k*_*m*_, respectively. The truncated PCE approximation of the solution reads

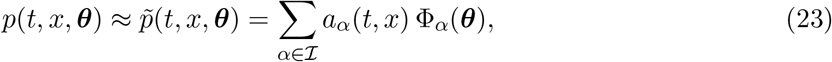

where *a*_*α*_ (*t, x*) are the deterministic PCE coefficients indexed by the multi-index *α* ∈ I. As in the one-parameter case, we substitute the PCE expansion into the PIDE and perform the projection step by multiplying by Φ_*k*_(***θ***) and taking the expectation with respect to the joint distribution of ***θ***. The resulting system of deterministic PIDEs for the coefficients *a*_*α*_ reads, for each *k* ∈ I,

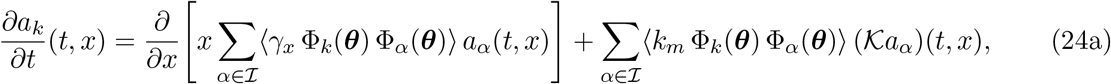

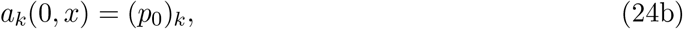

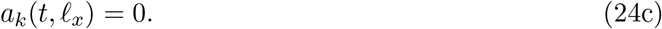

In contrast with the one-parameter case, the burst term now also introduces coupling between modes through the moments ⟨*k*_*m*_ Φ_*α*_Φ_*k*_⟩, since *k*_*m*_ is no longer deterministic. Defining the vector of PCE coefficients **a**^*L*^ := (*a*_*α*_)_*α∈I*_ the system can be written in vector form as

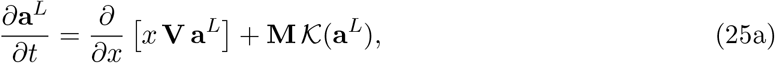

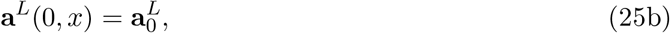

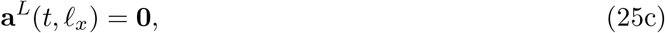

Where 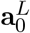 collects the PCE coefficients of the initial condition, and the matrices **V, M** ∈ ℝ ^*L×L*^ have entries

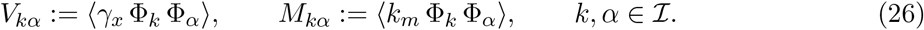

The previous procedure can be applied to any system and its complexity will depend on the nature of the original equations and the number of uncertain parameters.

### 2.4 Numerical Simulation

The resulting system of equations obtained from the projection steps is solved using a finite difference scheme for protein discretization. The protein number domain is discretized into a uniform grid consisting of *N* nodes with step size Δ*x* = *ℓ*_*x*_*/*(*N* − 1). The derivatives are approximated using summation by parts (SBP) operators [45] while boundary conditions are imposed using the simultaneous approximation term (SAT) method. Further details on the SBP-SAT framework can be found in [46], [47].

The SBP operator for the first derivative is constructed using a differentiation matrix **D** and a norm matrix **P**. For a generic grid function *w* with nodal values **w** = [*w*_1_, *w*_2_, …, *w*_*N*_ ]^*T*^ at the grid points *x*_*j*_, the first derivative is approximated as:

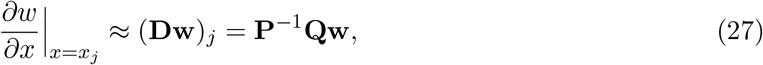

where **Q** satisfies the SBP property **Q** + **Q**^*T*^ = **E**_*N*_ − **E**_0_ = **B**, where **E**_*N*_ = diag(0, …, 0, 1) and **E**_0_ = diag(1, …, 0, 0) are diagonal matrices with entries corresponding to the boundary points of the domain [47].

Stacking the PCE coefficients **a**^*L*^(*t, x*_*j*_) at each of the *N* grid points into the global vector **a**(*t*) ∈ R^*NL*^ and applying the SBP-SAT discretization to the projected PIDE in (18) yields:

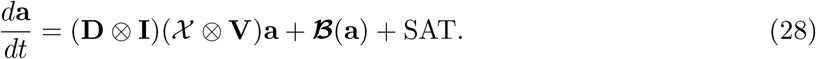

Here, X = diag(*x*_1_, *x*_2_, …, *x*_*N*_ ) is a diagonal matrix containing the protein grid points, ⊗; denotes the Kronecker product, and **I** is the *L* × *L* identity matrix acting on the PCE modes.

The SAT framework incorporates a term that enables the weak enforcement of boundary conditions, thereby preserving the stability of the numerical scheme [44]. Since the boundary condition is applied at the right border the SAT term is given by:

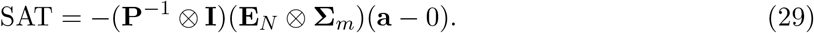

In this expression, **Σ**_*m*_ is a penalty matrix containing parameters that are chosen to ensure the stability of the numerical scheme. The derivation of the penalty parameter can be found in Annex A.

The time domain is discretized according to the Courant–Friedrichs–Lewy (CFL) restriction. Then, the time-step Δ*t* is chosen to satisfy the stability condition:

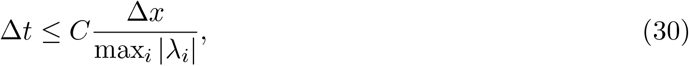

where *C* is the CFL number, Δ*x* is the protein grid size, and *λ*_*i*_ are the eigenvalues of the **V** matrix. Finally, the time integration is performed using an explicit fourth order Runge-Kutta method. The overall numerical scheme is designed to handle the coupled system of equations resulting from the PCE expansion while ensuring stability and convergence of the solution.

#### 2.4.1 Validation

The temporal domain considered is *t* ∈ [0, 20] and the protein number domain is *x* ∈ [0, 300]. The initial condition is taken as a standard Gaussian distribution. To assess convergence, we discretize the protein number domain using *N* ∈ {75, … 1500} grid points and consider PCE orders *d* ∈ {2, …, 6}.

For validation, we run Monte Carlo simulations with 10^4^ samples to produce reference solutions for the mean and variance of *p*(*t, x*) over time and protein number. Beyond these moments, the accuracy of the full reconstructed PDF is quantified using the total variation and Hellinger distances between the PCE surrogate and the reference solution. The former measures discrepancies in probability mass, while the latter captures differences in distributional shape through the squareroot densities [48]. For two probability densities on the protein domain [0, *ℓ*_*x*_], these distances are defined as

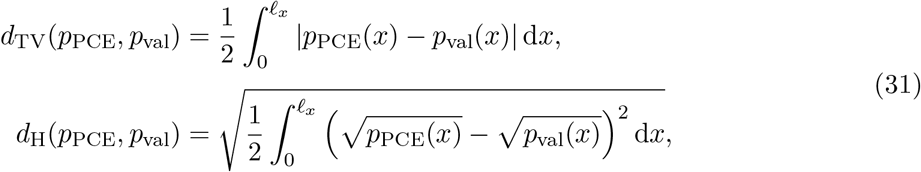

both taking values in [0, 1]. In addition, for the one-dimensional self-regulated case an analytical steady-state solution is available [22], [27], which provides an independent benchmark for the steadystate behavior. Reference solutions are computed using SELANSI, a MATLAB toolbox for fast evaluation of PIDEs [49]. The numerical and kinetic parameters used in these simulations are reported in Table 1.

**Table 1:** Kinetic and numerical parameter values used in the one-dimensional self-regulation PIDE simulations. Kinetic parameters were retrieved from [27], [49].

| Kinetic parameters |  |  |
| --- | --- | --- |
| Parameter | Description | Value |
| $H$ | Hill coefficient | 4 |
| $K$ | Hill constant | 45 |
| $\varepsilon$ | Leakage rate | 0.15 |
| $k_m$ | Transcription rate | $\mathcal{U}_{[8,10]}$ |
| $k_x$ | Translation rate | 400 |
| $\gamma_x$ | Protein degradation rate | $\mathcal{U}_{[0.8,1.2]}$ |
| $\gamma_m$ | mRNA degradation rate | 25 |
| Numerical simulation parameters |  |  |
| N | Grid points | 75–1500 |
| SBP order | Protein discretization | 4 |
| d | Polynomial chaos order | 2–6 |

## 3 Results

This section presents the results obtained from applying intrusive Galerkin projections to simulate one-dimensional PIDEs under parametric uncertainty. We explore two test cases: (i) a PIDE with an uncertain protein degradation rate, and, (ii) a PIDE with both uncertain degradation and transcription rates. The performance of the numerical schemes is evaluated in terms of accuracy, convergence, and computational efficiency.

### 3.1 One-dimensional PIDE with uncertain degradation rate

We first consider the self-regulation mechanism for a one-dimensional PIDE. In this example, protein degradation is not regulated by the protein level and enters the model as a linear term with rate *γ*_*x*_, whereas the probability of the promoter being in its active form depends on the amount of protein and is described by a Hill function. The binding of the protein to its own promoter constitutes a feedback regulation loop. The degradation rate *γ*_*x*_ is treated as an uncertain parameter, following a uniform distribution *γ*_*x*_ ∼ U_[0.8,1.2]_; accordingly, the polynomial basis functions are the normalized Legendre polynomials.

In the numerical simulations, fourth-order SBP operators are employed for protein discretization, and the time integration is performed using a fourth-order Runge-Kutta method. Figure 2a illustrates the convergence of the mean and variance of the number of proteins as the protein grid size and PCE order are refined. A minor effect on the surrogate error is observed with increasing order sizes of the PCE for the estimation of mean and variance. A substantial reduction in the *L*^2^ error over the protein domain occurs when refining the mesh from *N* = 75 to *N* = 300, followed by a plateau region for *N* ≥ 1200, where further refinement yields diminishing improvements. This behavior suggests that the stochastic truncation error is already spectrally converged for low PCE orders, and the dominant source of numerical error remains the protein discretization. The PCE surrogate also provides a flexible representation of the full protein PDF. Figure 2b shows arbitrary realizations of *p*(*x, γ*_*x*_) at *t* = 20 obtained directly from the precomputed PCE coefficients. The bimodal structure characteristic of cooperative positive feedback is captured across the entire range of *γ*_*x*_. Because the PCE coefficients are stored as time-resolved fields *a*_*j*_(*t, x*), this evaluation can be repeated at any time point of interest at negligible cost, providing a reconstruction of the protein PDF across both the parameter and the time dimensions.

**Figure 2:**
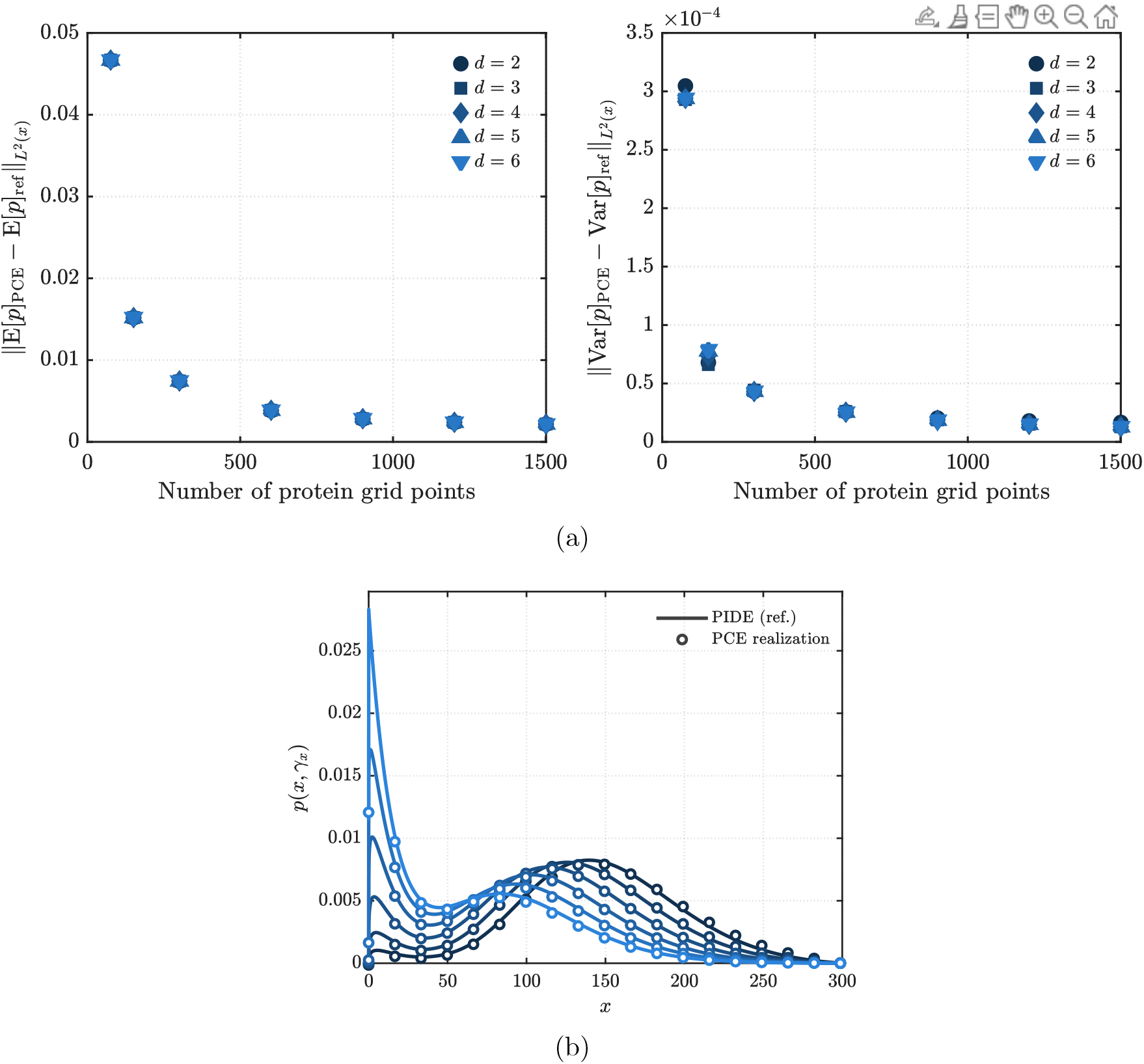
Results for the 1D PIDE with uncertain degradation rate. The results obtained from the intrusive PCE method are compared against Monte Carlo simulations with 10^4^ samples. Panel (a) shows the convergence of the mean and variance of the number of proteins as the protein grid is refined, for PCE orders ranging from *d* = 2 to *d* = 6. Panel (b) illustrates full protein PDF reconstructed for arbitrary realizations of the uncertain parameter at *t* = 20.

The accuracy of the PCE surrogate is further assessed by comparing its temporal evolution against the reference solution, as illustrated in Figure 3. The mean and variance of the number of proteins over time exhibit good agreement between the two methods. Note that, these two quantities of interest are obtained directly from the PCE coefficients using equations (12) without the need for additional sampling. This highlights one of the major advantages of the PCE approach: once the surrogate is built, the full statistical description of the system can be extracted at negligible computational cost. A complementary quantitative assessment of the full PDF reconstruction is reported in Figure 4, where the PCE surrogate is compared against the validation reference at *t* = 20 for the best-, median-, and worst-case realizations of *γ*_*x*_ selected by total variation distance and Hellinger distance. Across all parameter realizations, the total variation distance ranges from 2.5 × 10^*−*3^ to 3.17 × 10^*−*2^, while the Hellinger distance ranges from 3.8 × 10^*−*3^ to 3.66 × 10^*−*2^. The pointwise error *p*_PCE_ − *p*_val_ remains of order 10^*−*3^ across the three cases, with the largest deviations occurring in the region near *x* = 0.

**Figure 3:**
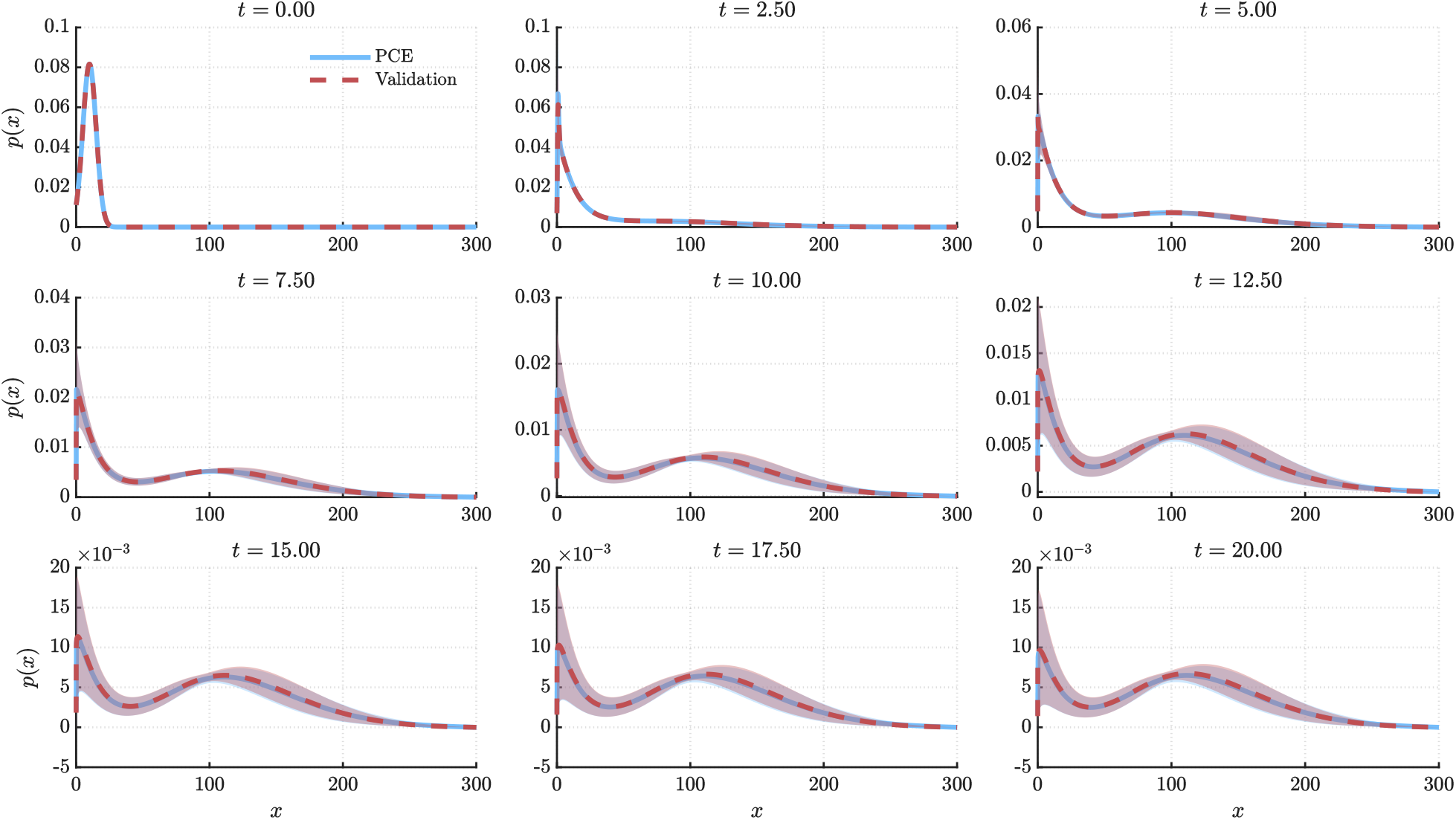
Time evolution of the protein probability density function *p*(*t, x*) predicted by the intrusive PCE method (solid blue) compared with the Monte Carlo reference solution based on 10^4^ samples (dashed red), for the one-dimensional PIDE with uncertain degradation rate. The shaded band depicts the spread induced by the uncertainty in *γ*_*x*_ (mean ± one standard deviation across realizations). Snapshots are reported at nine equispaced time points across the simulation horizon *t* ∈ [0, 20]. A PCE of order *d* = 2 and grid size *N* = 1500 was selected.

**Figure 4:**
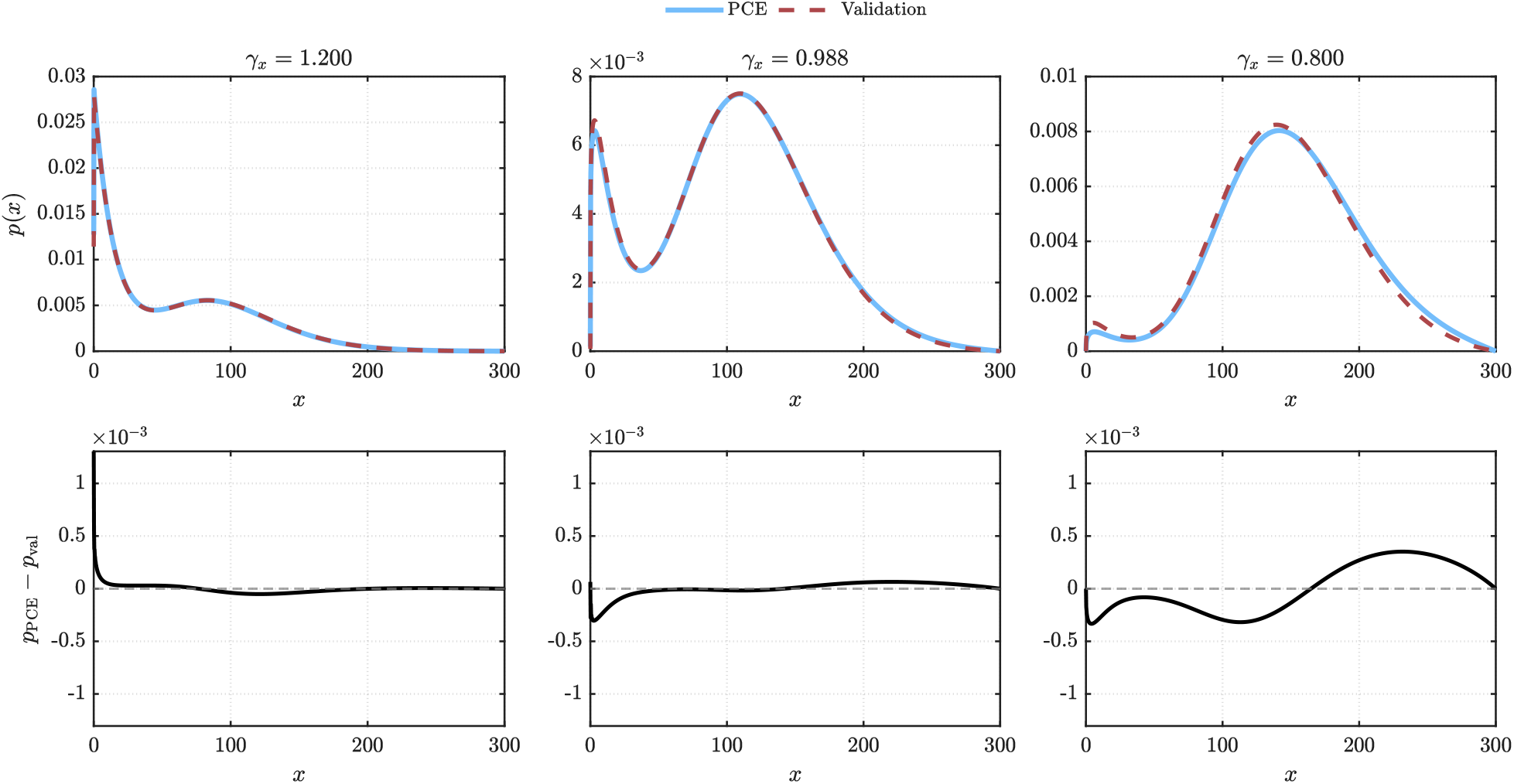
Comparison between the PCE surrogate and the validation reference at *t* = 20 for the best-, median-, and worst-case realizations of *γ*_*x*_ ranked by total variation distance. Top row: predicted PDFs (PCE solid blue, validation dashed red). Bottom row: pointwise error *p*_PCE_ − *p*_val_. PCE order *d* = 2 and grid size *N* = 1500.

A table summarizing the computational time for the different PCE orders and grid sizes is provided in Table 2. As expected, the computational time increases with both the PCE order and the grid size, since higher orders require more retained PCE coefficients and finer grids increase the cost of the underlying SBP discretization. In particular, the number of PCE coefficients grows combinatorially with the polynomial order, which directly increases the size of the coupled Galerkin system. Nevertheless, even in the most expensive configuration considered here (*N* = 1501, *d* = 6), the intrusive PCE solver completes in 8107 s (≈ 2.25 h). By contrast, generating the Monte Carlo validation set with 10^4^ samples using SELANSI requires approximately 10^4^ deterministic runs; with an average cost of about 2 s per run up to the defined simulation time, this corresponds to roughly 20,000 s (≈ 5.6 h). Importantly, once the PCE coefficients are computed, the solution statistics and the model response for new realizations of the uncertain parameters can be obtained by evaluating the polynomial expansion at negligible cost. This makes the approach particularly attractive for tasks that demand many repeated evaluations, such as optimization, parameter inference, and sensitivity analysis [34].

**Table 2:** Computational time in seconds for the intrusive PCE method applied to the onedimensional PIDE with uncertain degradation rate. The times are reported for variable PCE orders and grid sizes.

| Grid size (N) | $d = 2$ | $d = 3$ | $d = 4$ | $d = 5$ | $d = 6$ |
| --- | --- | --- | --- | --- | --- |
| 76 | 0.1475 | 0.1489 | 0.1681 | 0.2433 | 0.3885 |
| 151 | 0.4660 | 0.8280 | 1.304 | 2.053 | 2.999 |
| 301 | 3.721 | 9.728 | 35.58 | 42.00 | 68.65 |
| 601 | 92.58 | 170.0 | 281.2 | 398.7 | 511.0 |
| 901 | 305.8 | 573.2 | 746.2 | 946.4 | 1340 |
| 1201 | 453.1 | 913.0 | 1574 | 2323 | 3429 |
| 1501 | 911.4 | 1814 | 3080 | 4694 | 8107 |

### 3.2 One-dimensional PIDE with uncertain protein degradation rate and transcription rate

In this section, we extend the previous example by considering uncertainty in both the protein degradation rate *γ*_*x*_ and the mRNA transcription rate *k*_*m*_. Both parameters follow a uniform distribution, with degradation rate *γ*_*x*_ ∼ U_[0.8,1.2]_ and transcription rate *k*_*m*_ is modelled as *k*_*m*_ ∼ U_[8,10]_. Consequently, the polynomial basis functions employed are normalized Legendre polynomials. The temporal domain, initial conditions, and boundary conditions remain consistent with the previous example. Figure 5 illustrates the mean and variance of the number of proteins at time *t* = 20 obtained using a third-order PCE with a grid size of *N* = 1200. The results show that the intrusive PCE method accurately captures both the mean and the variance of the number of proteins, in good agreement with the reference solution from the validation set. The inclusion of uncertainty in both parameters leads to increased variability in the number of proteins, as reflected by the variance field.

**Figure 5:**
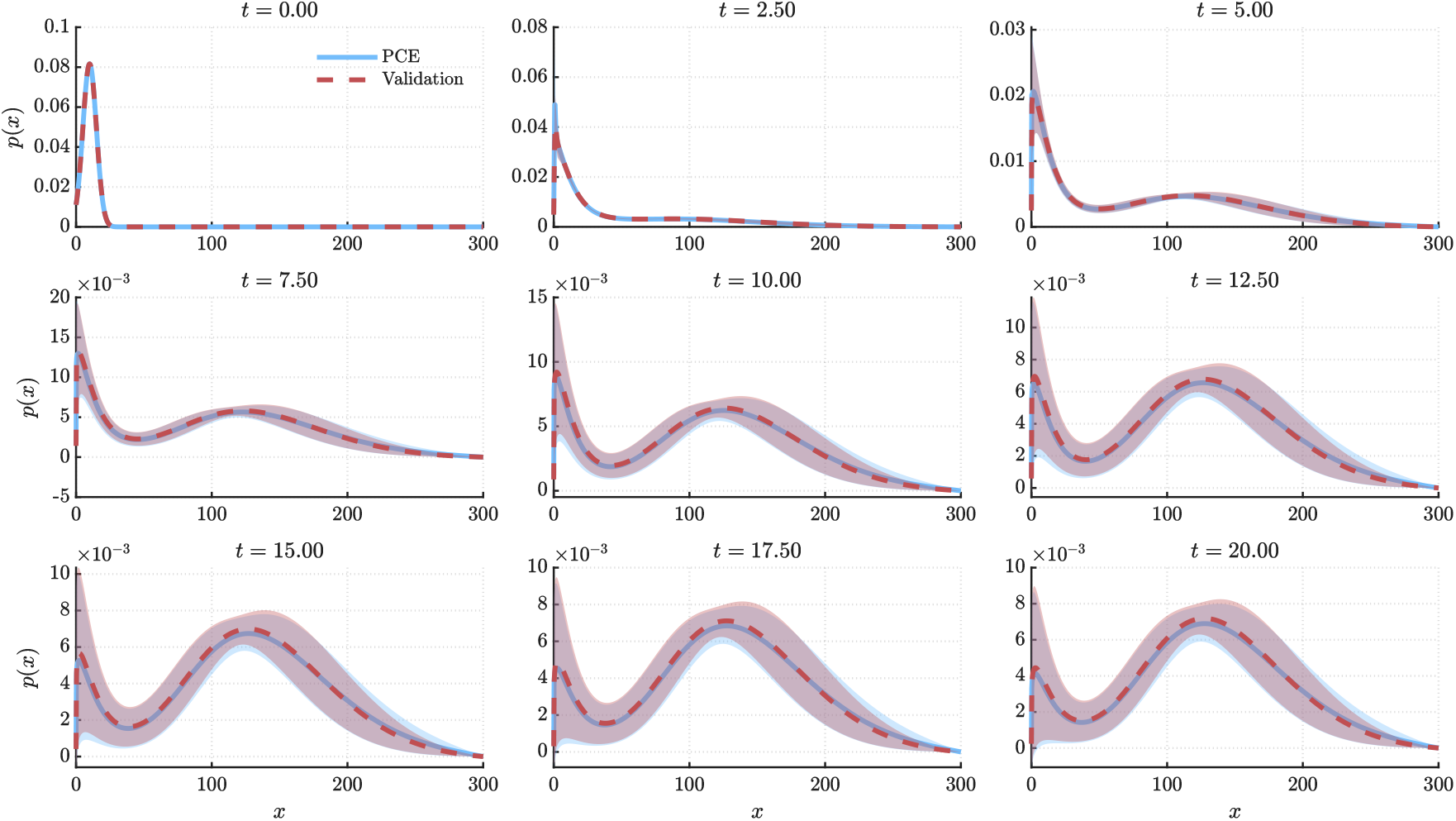
Time evolution of the protein probability density function *p*(*t, x*) predicted by the intrusive PCE method (solid blue) compared with the Monte Carlo reference solution based on 10^4^ samples (dashed red), for the one-dimensional PIDE with uncertain degradation rate and transcription rate. The shaded band depicts the spread induced by the uncertainty in *γ*_*x*_ (mean ± one standard deviation across realizations). Snapshots are reported at nine equispaced time points across the simulation horizon *t* ∈ [0, 20]. A PCE of order *d* = 3 and grid size *N* = 1200 was selected.

Nevertheless, the mean and variance are not sufficient to characterize the system behavior, because the protein distribution can undergo qualitative changes in shape. Figure 6 illustrates this effect as captured by the PCE surrogate: for low protein degradation rates *γ*_*x*_, the protein PDF can exhibit bimodality for specific values of *k*_*m*_. As *γ*_*x*_ increases, the bimodal structure progressively disappears and the distribution becomes unimodal, with the location of the transition depending on *k*_*m*_.

**Figure 6:**
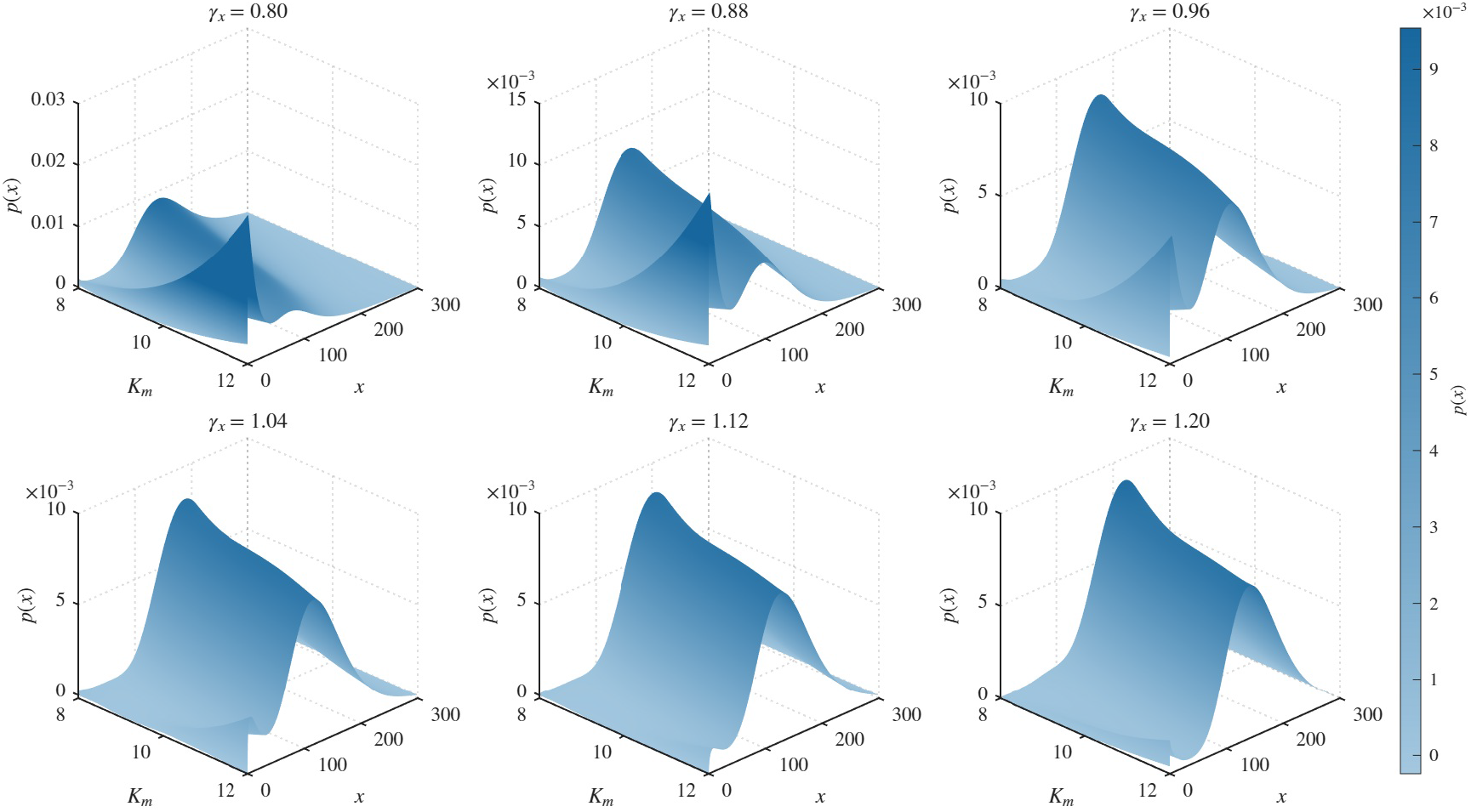
PCE distribution for the two-parameter uncertain PIDE. Different values for *k*_*m*_ are considered while simulating the solutions for variable degradation rates at *t* = 20. Solutions are obtained with PCE of order three for each parameter and grid size *N* = 1200.

In contrast to the single-parameter case, where the convergence analysis showed that the PCE order had only a minor effect on the surrogate accuracy, the two-parameter setting required increasing the polynomial order to *d* = 3 to achieve a comparable level of agreement on the full PDF. Figure 7 shows the best-, median-, and worst-case total variation and Hellinger distances at steady state for this PCE configuration with grid size *N* = 1200. The need for higher-order expansions is particularly pronounced in parameter regimes where the distribution transitions between unimodality and bimodality.

**Figure 7:**
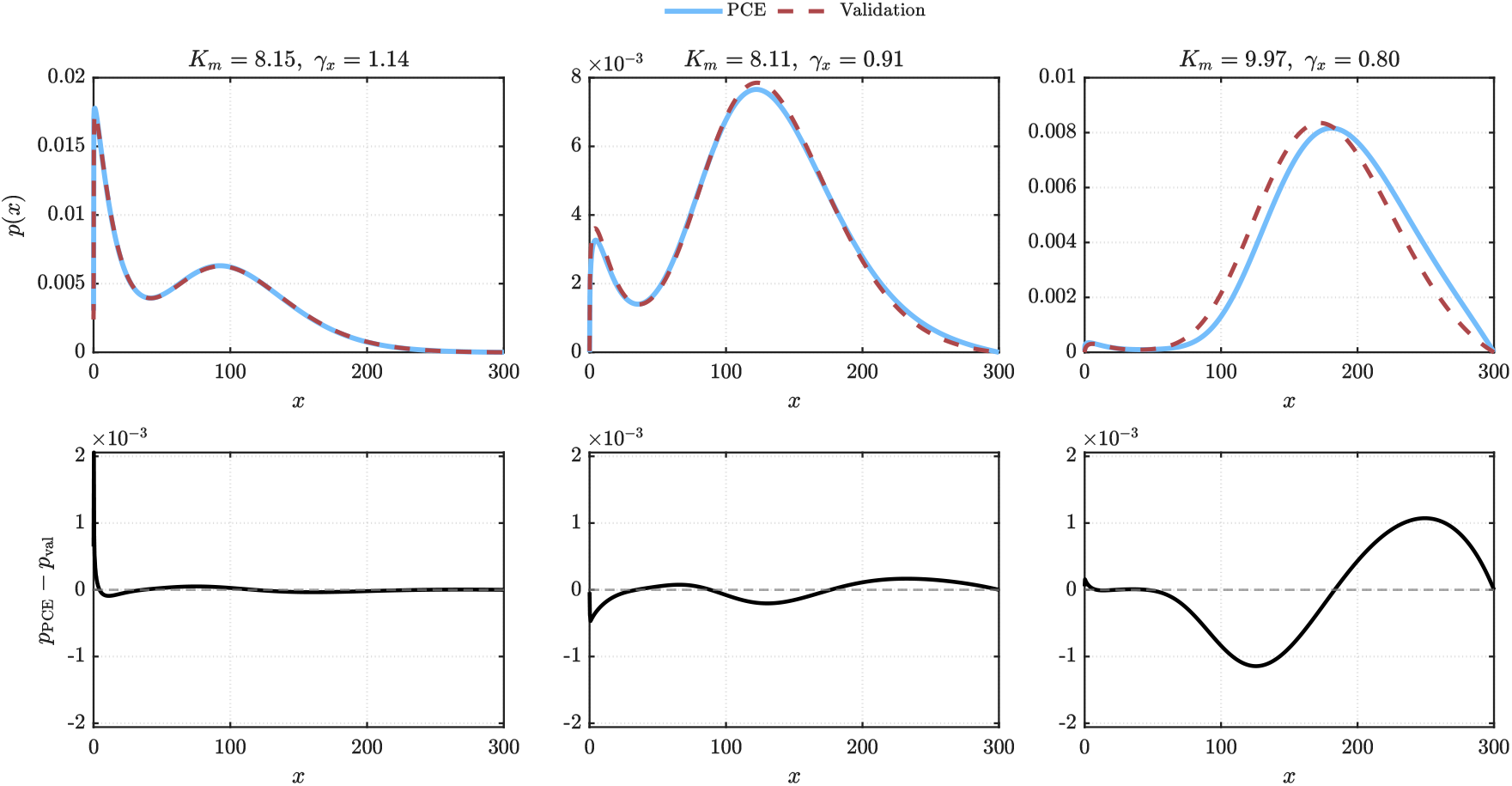
Comparison between the intrusive PCE surrogate and the reference validation solution at time *t* = 20 for a third order expansion and grid size *N* = 1200. The top row shows the probability density functions of the number of proteins, while the bottom row displays the corresponding pointwise error *p*_PCE_ − *p*_val_. The three columns correspond to the best-, median-, and worst-case realizations selected according to the total variation distance. Across all realizations, the total variation distance ranges from 2.4 × 10^*−*3^ to 8.29 × 10^*−*2^, while the Hellinger distance ranges from 4.7 × 10^*−*3^ to 8.20 × 10^*−*2^.

Nevertheless, the mean and variance are not sufficient to characterize the system behavior, because the protein distribution can undergo qualitative changes in shape. Figure 6 illustrates this effect as captured by the PCE surrogate: for low protein degradation rates *γ*_*x*_, the protein PDF can exhibit bimodality for specific values of *k*_*m*_. As *γ*_*x*_ increases, the bimodal structure progressively disappears and the distribution becomes unimodal, with the location of the transition depending on *k*_*m*_.

In contrast to the single-parameter case, where the convergence analysis showed that the PCE order had only a minor effect on the surrogate accuracy, the two-parameter setting required increasing the polynomial order to *d* = 3 to achieve a comparable level of agreement on the full PDF. Figure 7 shows the best-, median-, and worst-case total variation and Hellinger distances at steady state for this PCE configuration with grid size *N* = 1200. The need for higher-order expansions is particularly pronounced in parameter regimes where the distribution transitions between unimodality and bimodality.

The best and median cases correspond to parameter regimes in which bimodality is observed, where the surrogate captures both modes with good agreement. In contrast, the worst-case error occurs in regimes where the distribution is strictly unimodal and the solution deviates most strongly from the mean behavior of the system. This deterioration is consistent with a well-known limitation of global PCE representations: when the map from uncertain inputs to outputs becomes highly non-smooth, convergence can slow down or fail [50], [51].

## 4 Conclusion

In this work, we developed a framework for uncertainty quantification in stochastic gene regulatory networks by combining PIDE models with intrusive polynomial chaos expansions and Galerkin projections. The PIDE formulation captures the time evolution of protein distributions under intrinsic biochemical noise, while parametric uncertainty is introduced through uncertain kinetic rates to represent extrinsic variability. Projecting the resulting PIDE onto an appropriate polynomial chaos basis yields a coupled deterministic system whose solution provides direct access to statistics of interest and to approximations of the full probability density function. The approach was assessed on a one-dimensional gene expression model with uncertain degradation and transcription rates, where it accurately reproduced key distributional features, including regime-dependent transitions between unimodal and bimodal behavior. Once computed, the PCE surrogate provides rapid predictions for new parameter realizations by simple evaluation of the expansion, avoiding repeated PIDE solves. This makes uncertainty-aware inference, optimization, and control practical for stochastic GRN models, where Monte Carlo–based approaches would be prohibitively expensive.

Several limitations should be acknowledged. The size of the projected Galerkin system grows combinatorially with both the number of uncertain parameters and the polynomial order, which constrains the present formulation to low- and moderate-dimensional uncertainty; sparse and adaptive PCE constructions are natural avenues to mitigate this scaling. In addition, the validation is restricted to one-dimensional self-regulatory networks under continuous and independent parametric distributions, and the extension to multi-protein cross-regulatory networks, correlated parameters, and discrete uncertainty sources will require further methodological development.

Based on these observations, future work will focus on scalable solvers for the resulting coupled Galerkin systems and on extending the formulation to richer regulatory architectures. Possible applications of the resulting surrogate include uncertainty-aware parameter inference and optimization, as well as control tasks such as efficient PIDE-based model predictive control (MPC), as recently explored in [52].

## A Derivation of the penalty terms

In this annex, we derive the penalty parameters used in the simultaneous approximation term (SAT) by means of an energy stability analysis.

We consider the projected system

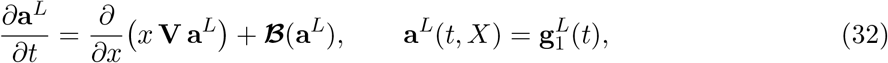

where **V** = **V**^*⊤*^ is the stochastic Galerkin advection matrix. We derive the SAT penalty parameter by an energy method and focus on the transport term and the boundary treatment. For brevity, in what follows we drop the superscript *L* and write **a** := **a**^*L*^.

We define the discrete energy as

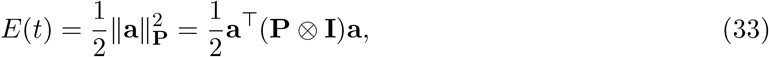

where **P** is the symmetric positive definite norm matrix associated with the SBP operator, **I** is the identity matrix in the stochastic dimension, and ⊗ denotes the Kronecker product.

Taking the time derivative of the energy yields

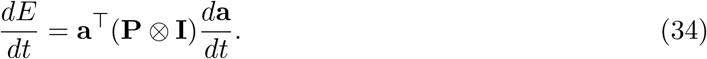

Substituting the semi-discrete form of the PIDE (28) and retaining only the transport and boundary penalty terms, we obtain:

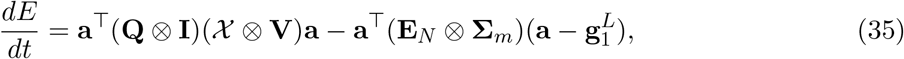

where **Q** = **PD** is the SBP differentiation matrix and X = diag(*x*_1_, …, *x*_*N*_ ).

Using the SBP property **Q** + **Q**^*⊤*^ = **B** := **E**_*N*_ − **E**_0_ together with the symmetry of **V**, the transport contribution reduces to boundary terms. Indeed,

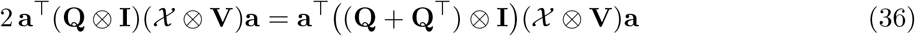

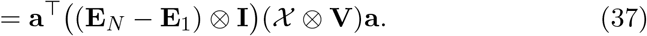

Hence,

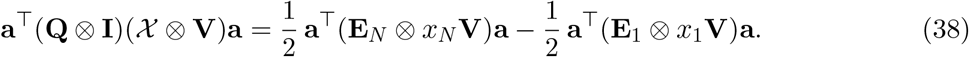

Since the left boundary does not contribute (e.g. *x*_1_ = 0), we obtain

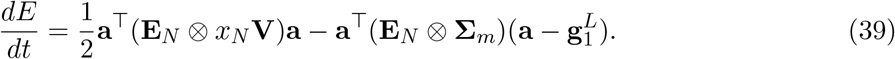

In particular, for homogeneous boundary data 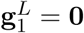 and collecting the terms associated with the right boundary node yields

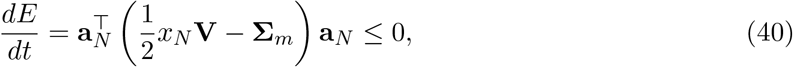

where **a**_*N*_ denotes the vector of stochastic coefficients at the boundary node *x* = *x*_*N*_ . For energy stability, we require 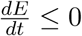, which is guaranteed if

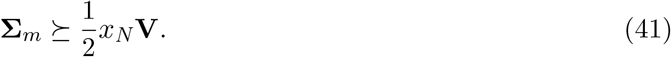

